# Noise Effect on the Temporal Patterns of Neural Synchrony

**DOI:** 10.1101/2021.03.22.436529

**Authors:** Joel Zirkle, Leonid L Rubchinsky

## Abstract

Neural synchrony in the brain is often present in an intermittent fashion, i.e., there are intervals of synchronized activity interspersed with intervals of desynchronized activity. A series of experimental studies showed that this kind of temporal patterning of neural synchronization may be very specific and may be correlated with behavior (even if the average synchrony strength is not changed). Prior studies showed that a network with many short desynchronized intervals may be functionally different from a network with few long desynchronized intervals as it may be more sensitive to synchronizing input signals. In this study, we investigated the effect of channel noise on the temporal patterns of neural synchronization. We employed a small network of conductance-based model neurons that were mutually connected via excitatory synapses. The resulting dynamics of the network was studied using the same time-series analysis methods as used in prior experimental and computational studies. While it is well known that synchrony strength generally degrades with noise, we found that noise also affects the temporal patterning of synchrony. Noise, at a sufficient intensity (yet too weak to substantially affect synchrony strength), promotes dynamics with predominantly short (although potentially very numerous) desynchronizations. Thus, channel noise may be one of the mechanisms contributing to the short desynchronization dynamics observed in multiple experimental studies.

**Highlights:**

- Channel noise alters the temporal pattern of intermittent neural synchrony
- Noise may alter this pattern without significant change in average synchrony strength
- The resulting patterning is similar to that observed in multiple experiments

## 1. INTRODUCTION

Neural systems exhibit synchronization of oscillatory activity in a variety of different situations. This synchronization is involved in multiple brain functions in cognitive and motor domains (for example, see reviews by Buzsáki and Draguhn, 2004; Fell and Axmacher, 2011; Fries, 2015; Harris and Gordon, 2015). Disorganization of neural synchrony (such as excessively strong or insufficiently strong synchrony) negatively affects the information processing in the networks of the brain and is associated with several neurological disorders (such as Parkinson’s disease) and psychiatric disorders (such as schizophrenia and autism spectrum disorders), see (Schnitzler and Gross, 2005; Uhlhaas and Singer, 2006; Oswal et al., 2013; Pittman-Polletta et al., 2015).

Perfect synchrony is probably not achievable in the brain (at least in the rest state). A moderate synchrony strength implies that synchrony is sometimes high and sometimes low, yielding some average synchrony strength level. This kind of temporal patterning of synchronous activity may be independent of the synchronization strength (Ahn et al., 2018). Time-series analysis techniques to quantify the temporal patterning of synchrony on very short time-scales (provided there is a statistically significant synchrony overall) have been developed over the past decade (e.g., Park et al., 2010; Ahn et al., 2011). These techniques revealed that neural synchrony in the brain shows a very specific patterning: it is interrupted by potentially numerous but very short desynchronization episodes. This was observed in different species (rodents, humans), different brain signals (spikes, LFPs, EEG), different brain areas (cortex, hippocampus, basal ganglia), and different brain states (healthy and diseased), see (Park et al., 2010; Ahn and Rubchinsky, 2013; Ahn et al., 2014; Ratnadurai-Giridharan et al., 2016; Malaia et al., 2020) for the different experiments. The distribution of desynchronization durations is altered under different conditions. It was found to be related to the severity of symptoms of Parkinson’s disease, addiction, and autism (Ahn et al., 2014; Ahn et al., 2018; Malaia et al., 2020, Dos Santos Lima et al., 2020). However, the mode of this distribution is always one (see studies mentioned above).

Thus, the mechanisms behind the short desynchronization dynamics make an important problem to explore. Kinetics of sodium and potassium spike-producing ionic channels in neurons as well as spike-timing dependent plasticity can facilitate short desynchronization dynamics (Ahn and Rubchinsky, 2017; Zirkle and Rubchinsky, 2020). Noise can also potentially be a factor in temporal patterning of synchrony, because noise is well-known to affect synchronous dynamics in multiple ways from a straightforward decrease of synchrony under the action of noise to the increase of synchrony due to correlated noisy input (Zhou et al., 2002; Goldobin and Pikovsky, 2005) and can exert multiple effects on the synchrony between and within networks of neurons (e.g., McMillen and Kopell, 2003; Zhou et al., 2013; Meng and Riecke, 2018).

Noise is ubiquitous in neural systems (Ermentrout et al., 2008, Faisal et al., 2008). Thus, the question of how noise may affect the temporal patterning of neural synchrony is very natural. Leaving aside the issue of what is noise in neural systems (Stein et al., 2005; Yarom and Hounsgaard, 2011), we will focus here on specific types of noise. We will consider the effect of channel noise in individual neurons (multiplicative noise) as well as the effect of an additive noise on the temporal patterning of synchronized dynamics in a network of synaptically coupled model neurons. We show that both noise types can robustly alter the temporal patterning of the synchronized dynamics, effectively shortening desynchronization intervals while simultaneously making them more frequent. This is similar to what one can observe in experimental data.

## 2. METHODS

To focus on the very basic aspects of the noise effect on the temporal patterns of synchronization, we employ a minimal heterogeneous network of relatively simple model neurons following (Ahn and Rubchinsky, 2017).

### 2.1. Neuronal and Synaptic Modeling

We use the same neuronal and synaptic model as the one used in (Ahn and Rubchinsky, 2017) and incorporate noise into it. This is a two-dimensional ODE model, a simplification of the Hodgkin-Huxley model which is equivalent to the Morris-Lecar model. It is one of the simplest models of neurons that retain kinetics of ionic channels (which may be important for the temporal patterns of synchronized dynamics as suggested by (Ahn and Rubchinsky, 2017)). The model takes the following form:

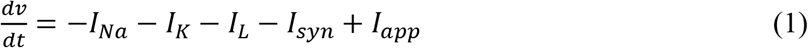

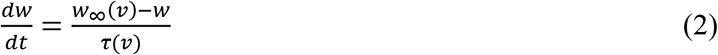

Here *v* is the membrane potential of a neuron and *w* is the gating variable for the potassium current. The synaptic current between neurons, *I*_*syn*_, is described below. *I*_*app*_ is a constant input current to each neuron which controls the excitability of the cell. The spike-producing sodium and potassium currents and the leak current are described by:

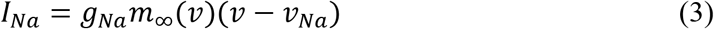

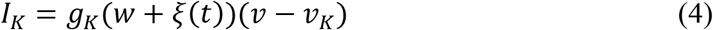

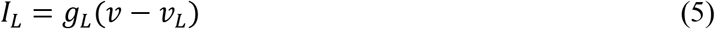

*g*_*Na*_, *g*_*K*_, *g*_*L*_ are the maximal conductances for the sodium, potassium and leak currents, respectively. A Gaussian white noise term, *ξ*(*t*), is added to the gating variable *w*. This introduction of noise represents the inherently stochastic nature of the membrane ion channels in neurons (Goldwyn and Shea-Brown, 2011). The steady-state values for the gating variables of the sodium and potassium currents are:

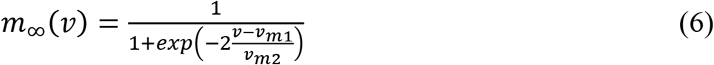

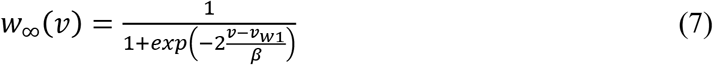

The voltage-dependent activation time constant of the potassium current is:

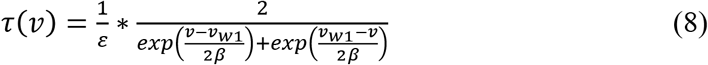

All synapses are excitatory, and the synaptic current from each neuron *j* ≠ *i* to neuron *i* is given by:

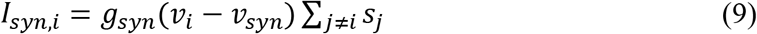

*g*_*syn*_ is the maximal conductance of the synapse (i.e., the synaptic strength), and *s*_*j*_ is the synaptic variable for neuron *j* and the summation is taken over all neurons that are connected to the *i*-th neuron. The synaptic variable *s* is governed by:

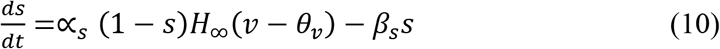

*H*_∞_ is a sigmoidal function whose input is the presynaptic neuronal voltage:

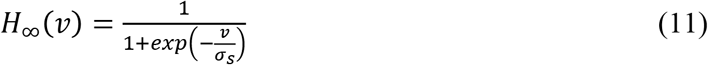

With the exception of ∝_*s*_, the values of the cellular and synaptic parameters are the same as used in (Ahn and Rubchinsky, 2017): *g*_*Na*_ = 1, *g*_*K*_ = 3.1, *g*_*L*_ = 0.5, *v*_*Na*_ = 1, *v*_*K*_ = −0.7, *v*_*L*_ = −0.4, *v*_*m*1_ = −0.01, *v*_*m*2_ = 0.15, *v*_*w*1_ = 0.08, *β* = 0.145, *I*_*app*_ = 0.045, *ε*_1_ = 0.02 (unless noted otherwise as in 3.1 where different values of *ε* are considered) and *ε*_2_ = 1.2*ε*_1_ are values of *ε* in two different neurons, *v*_*syn*_ = 0.5, ∝_*s*_ = 2, *β*_*S*_ = 0.2, *θ*_*v*_ = 0.0, *σ*_*s*_ = 0.2.

We also simulated the neurons with a current noise (additive noise) instead of a channel noise. The model is identical to the one described above, except that the noise term is now an additive term in the voltage equation:

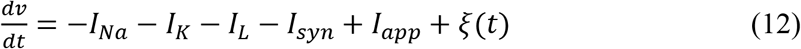

### 2.2. Numerical Implementation

Multiplying out the voltage equation from the multiplicative noise section, we obtain the following Langevin-type equation:

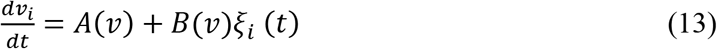

where *A*(*v*) and *B*(*v*) are the drift and diffusion terms, respectively. In the case of additive noise we simply have

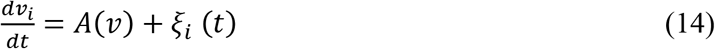

We then solve the system numerically using the Euler-Maruyama method (Higham, 2001; Gardiner, 2009). Here *ξ*(*t*) is white noise that is distributed as 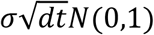, where *σ* ∈ [0,0.02] is the noise strength. This interval was chosen so that the noise could be strong enough to induce a change in the temporal patterns of synchronous dynamics, yet not destroy the inherent spiking dynamics. The noise term for each neuron is generated with a different seed, i.e., the noise terms for each neuron are uncorrelated. The unit of time in our system is millisecond. The system was integrated on the time interval [0,20000] with a time step of *dt* = 0.01 ms. To account for the initial transient behavior the first 5% of the time-series was discarded from analysis.

Depending on which parameter value was varied, the voltage threshold to define an action potential was set at either 0.20, 0.25 or 0.30. To eliminate the possibility of the channel noise driving the membrane voltage over the threshold immediately following an action potential, a window of 15 ms was set after each neuron’s action potential in which we do not count threshold crossings. Since the highest recorded frequency was approximately 40 Hz, a window of 15 ms is appropriate.

### 2.3. Synchronization Data Analysis

The time-series analysis of synchronized dynamics in the network follows the earlier study of a non-noisy version of this model (Ahn and Rubchinsky, 2017), and is similar to the analysis of the temporal patterns of neural synchrony in the experimental studies mentioned in the Introduction. We will briefly describe the major steps of the analysis here.

The phase, *φ*(*t*), of a neuron is defined as

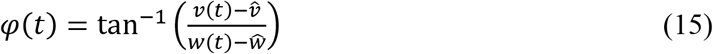

where (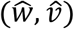) is a point selected inside the neuron’s limit cycle in the (*w*, *v*) – plane. The synchronization strength is computed as

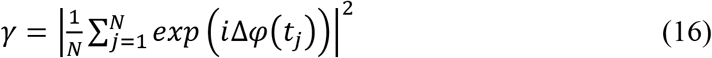

where Δ*φ*(*t*_*j*_) = *φ*_1_(*t*_*j*_) − *φ*_2_(*t*_*j*_) is the difference of the phases of neurons 1 and 2 at time *t*_*j*_. *N* is the number of data points. The value of *γ* ranges from 0 to 1, which represent a complete lack of synchrony and perfect synchrony, respectively.

The index *γ* represents an average value of phase-locking over the interval of analysis. However, to describe how synchrony varies in time, one needs to look at the transitions to and from a synchronized state on short timescales. This is done as follows.

If there is some degree of phase-locking present, then there is a synchronized state, i.e., a preferred value of the phase difference Δ*φ*. We check if the actual phase difference is close to this preferred value or not for each cycle of oscillations. When *φ*_1_ increases past zero, say at time *t*_*i*_, then *φ*_2_(*t*_*i*_) is recorded. This generates a sequence of numbers 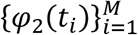 Due to the presence of some synchrony, there is a clustering about some phase value, say *φ*_0_. This is taken as the preferred phase value, and if *φ*_2_(*t*_*i*_) = *φ*_*i*_ differs from it by more than a threshold value of 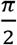 (the same value as in the experimental studies described in the Introduction) then the neurons are considered to be in the desynchronized state, otherwise they are considered to be in the synchronized state.

The length of a desynchronization event is defined as the number of cycles the system spends in a desynchronized state minus one. The lengths of all desynchronization events are recorded, and the mode of these lengths is considered as a way to describe the distribution of desynchronizations (as in the experimental studies). For later reference, a “mode *n*” system or “mode *n*” dynamics means that the mode of all lengths of desynchronization events for that particular system (that particular set of parameter values) is *n*. Note that in this approach, the duration of desynchronizations is measured in relative units (the number of cycles of oscillations). We present examples of time-series and associated distributions of desynchronization durations below in the beginning of the Results section.

## 3. RESULTS

Following (Ahn and Rubchinsky, 2017), we used a simple heterogeneous network consisting of two neurons connected via excitatory synapses with the same synaptic strength (see Fig. 1). As described in Methods, the values of ε (determining the potassium current activation dynamics) differ slightly between the two neurons, *ε*_2_ = 1.2*ε*_1_, hence their frequency of spiking is slightly different. Also, the strength of both synapses is *g*_*syn*_ = 0.005, hence the coupling is weak. Due to the heterogeneity of the network and the weak coupling, the synchronization strength in the network is relatively weak. While this study considered network heterogeneity due to different values of ε in neurons, we also performed a series of simulations, where a similar level of heterogeneity is achieved via difference of *g*_*K*_ or *I*_*app*_. The resulting effect of the noise action on the patterns of synchrony was similar to the one observed in the networks heterogeneous in ε, and thus was not studied in detail and is not reported here.

**Figure 1.**
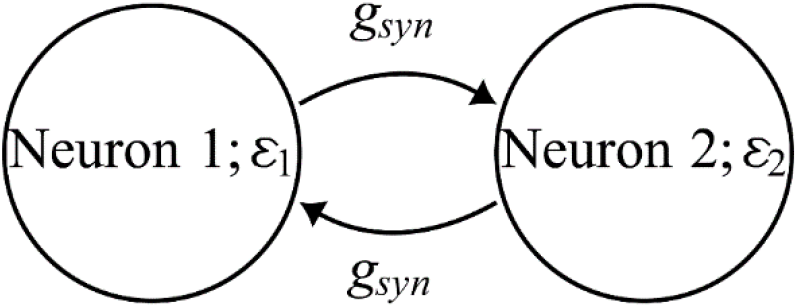
The diagram of the minimal network: two cells mutually connected with mutually excitatory synapses.

Figure 2 presents an example of time-series of two neurons (Fig. 2A) and phase plane illustrations of these neurons (Fig. 2B) in uncoupled, coupled, and coupled with noise cases. Note that the coupling is not strong enough to lead to strong synchrony, and that noise levels considered here do not substantially alter the shape of the spike and phase plane trajectory or substantially affect synchrony strength. This lets us to focus on the action of noise on the temporal patterning of synchronous dynamics.

**Figure 2.**
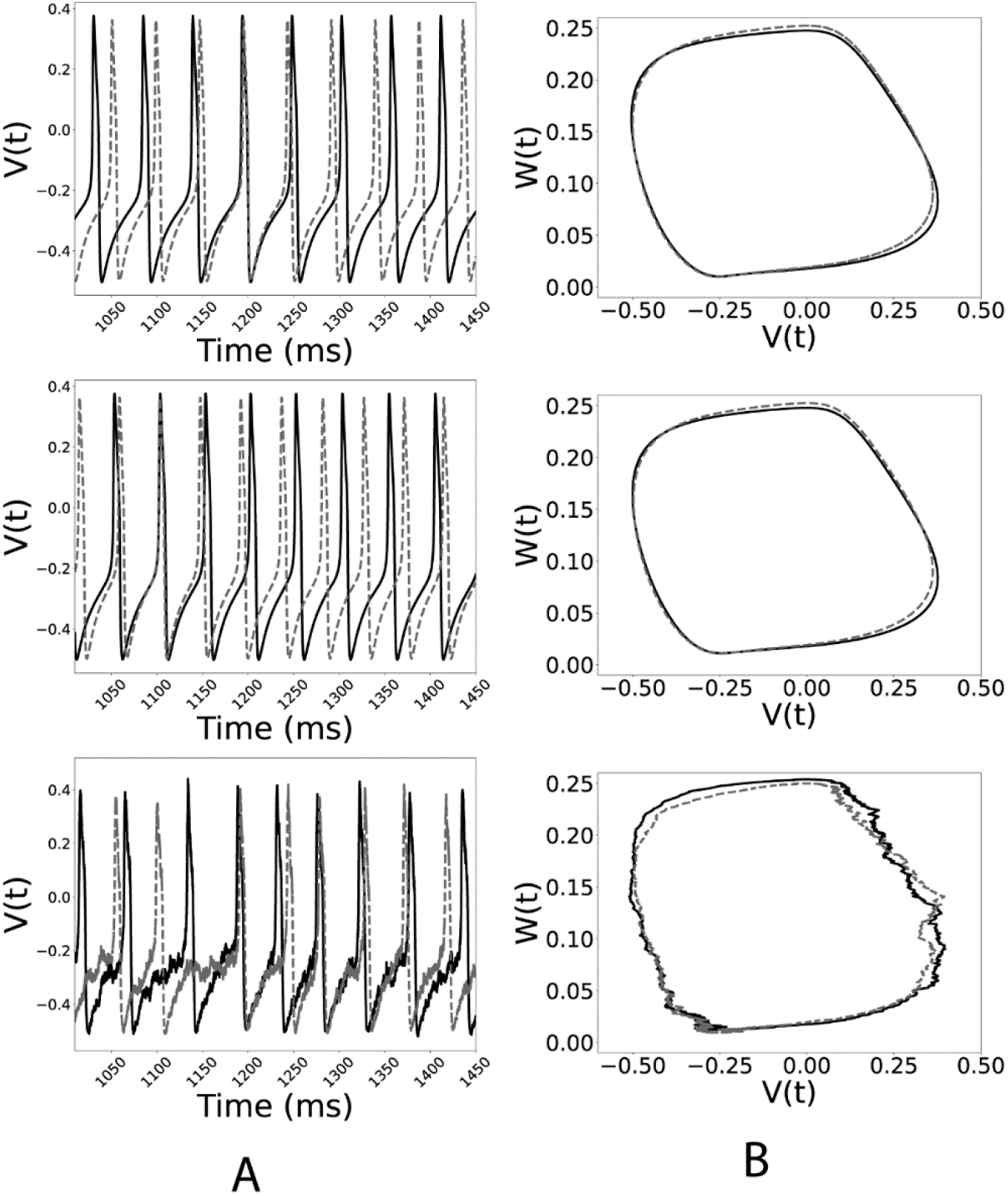
Voltage time-series (A) and phase plane trajectories (B) of two neurons in the network (solid black and dashed grey lines). Top row is the case of uncoupled neurons, middle row is the case of weakly coupled neurons, and the bottom row is the case of coupled neurons with noise (*ε*_1_ = 0.044 and the noise level in the bottom row is *σ* = 0.0149).

Based on the noiseless case considered in (Ahn and Rubchinsky, 2017) and similarly to (Zirkle and Rubchinsky, 2020), we vary the values of parameters of the potassium current kinetics in such a way as to change the dynamics of the noiseless network from exhibiting predominantly short desynchronizations (i.e., those observed in experiments) to one with longer desynchronizations. These parameters are *ε*; the reciprocal of the peak value of the activation time-constant *τ*(*v*), *β*; the widths of the activation time-constant *τ*(*v*) and the steady-state activation function *w*_∞_(*v*), and *v*_*w*1_; the voltage of half-activation and of the peak activation constant. Variation of either of these three parameters effectively changes the voltage-dependent activation time-constant *τ*(*v*) to either large or small, which delays or accelerates the activation of potassium current, respectively (and changes the waveform of oscillations from sharp to more sinusoidal).

### 3.1. Variation of *ε*

We first will explore if the channel noise preserves the prevalence of short desynchronizations or not. Smaller values of *ε* correspond to larger values of τ(*v*) and are known to promote short desynchronization dynamics (Ahn and Rubchinsky, 2017); smaller values of *ε* also result in a lower frequency. For *ε*_1_ = 0.044, the noiseless system is mode 1 (i.e., the mode of the distribution of all desynchronization event lengths is one cycle of oscillation). This is similar to the experimental results (see Introduction). As the strength of the noise is increased, we see from Fig. 3A that the system remains mode 1. It is worth noting, that the average synchrony strength stays virtually the same (Fig. 3B). Of course, a very strong noise will change it, but the noise range we consider here is sufficiently weak so as to not affect the average synchrony strength γ. We also plot the frequency of oscillations (firing rate) in Fig. 3C. The noise does not change it in a substantial way either, although we include it here because it is affected by the change of *ε* and other channel parameters.

**Figure 3.**
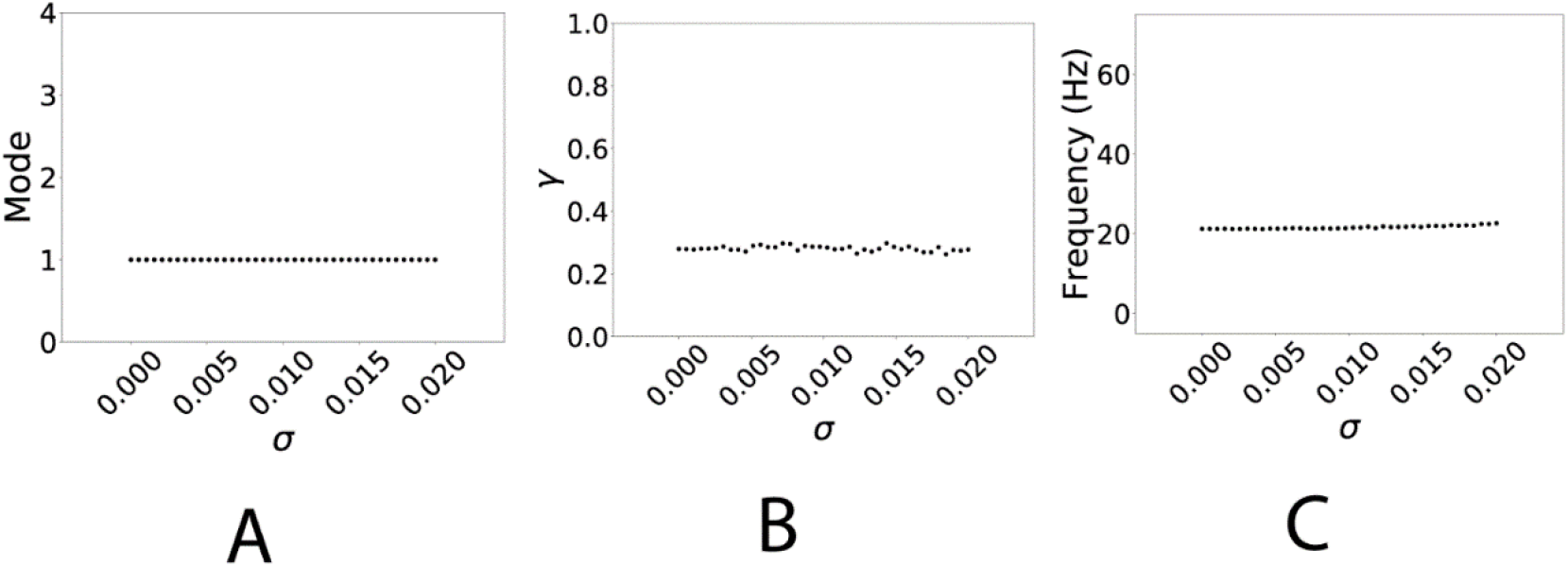
The effect of noise on synchrony properties of the network for the case of small *ε* (*ε*_1_ = 0.044), which exhibits mode 1 desynchronization dynamics in the noiseless case. The strength of the noise σ is varied along the horizontal axes. A: mode of the distribution of desynchronization durations, B: average synchrony index *γ*, C: mean oscillation frequency (firing rate) of the system.

The effect of channel noise on the distribution of desynchronization durations for *ε*_1_ = 0.044 is shown in Figure 4. The strength of the noise increases from left to right, and the effect is to broaden the distribution while maintaining a mode of 1. Fig. 4C for instance, is qualitatively similar to distributions obtained from previous studies conducted on experimental data (see references in Introduction).

**Figure 4.**
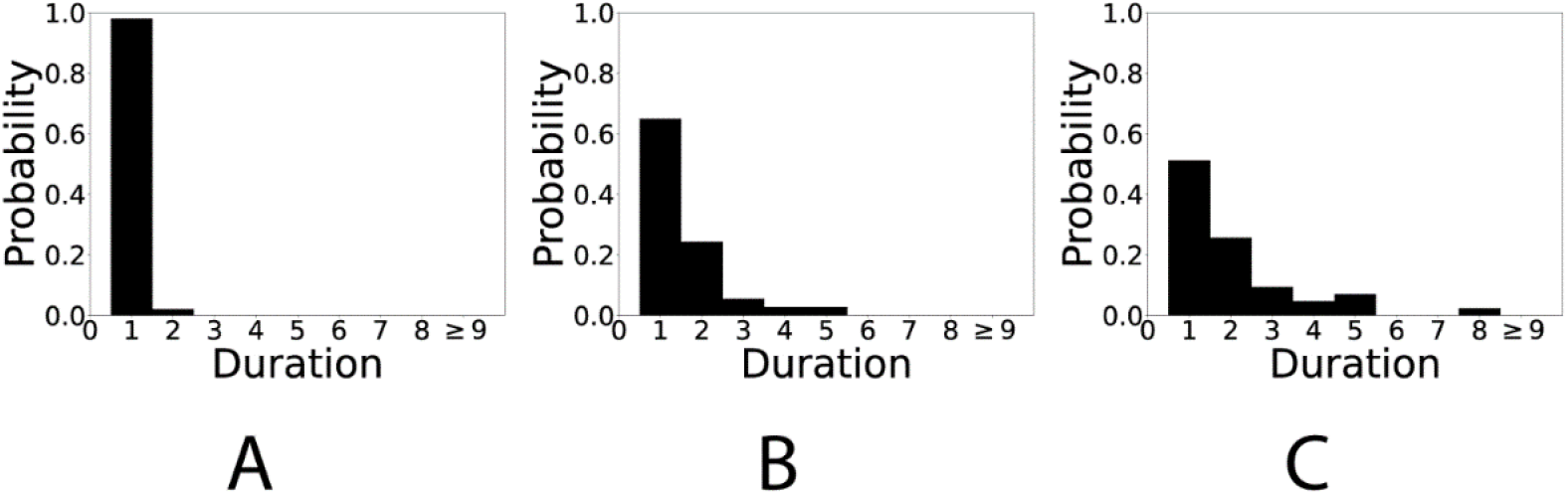
Distributions of desynchronization durations for different levels of noise in the network exhibiting mode 1 desynchronization dynamics in the noiseless case (*ε*_1_ = 0.044). Noise strength increases from left to right: A: *σ* = 0 (no noise), B: *σ* = 0.0097, C: *σ* = 0.02.

We now explore how the network with longer desynchronizations in the noiseless case will respond to noise. For this we consider a higher value of *ε*; for *ε*_1_ = 0.132, the noiseless network is mode 2. As the noise strength is increased the mode of the system shifts from 2 down to 1 (Fig. 5A). This is not a fully monotonous transition. Several other noise-induced transitions between dynamics with different modes considered below are not monotonous either. Moreover, the mode is just a convenient characteristic of the desynchronization episodes. They may experience more gradual changes not captured by the mode. However, the results indicate that on a larger scale, there is a clear noise-induced transition from longer to shorter desynchronizations. Once again, the average synchrony strength and the frequency of firing are nearly constant with respect to the noise strength (Fig. 5B and 5C). This will be the case for all the situations considered here, reflecting the fact that the noise we use is effectively weak. This indicates that the distribution of the desynchronization durations can be altered independently of the average synchrony strength and the frequency of oscillations by varying the noise strength.

**Figure 5.**
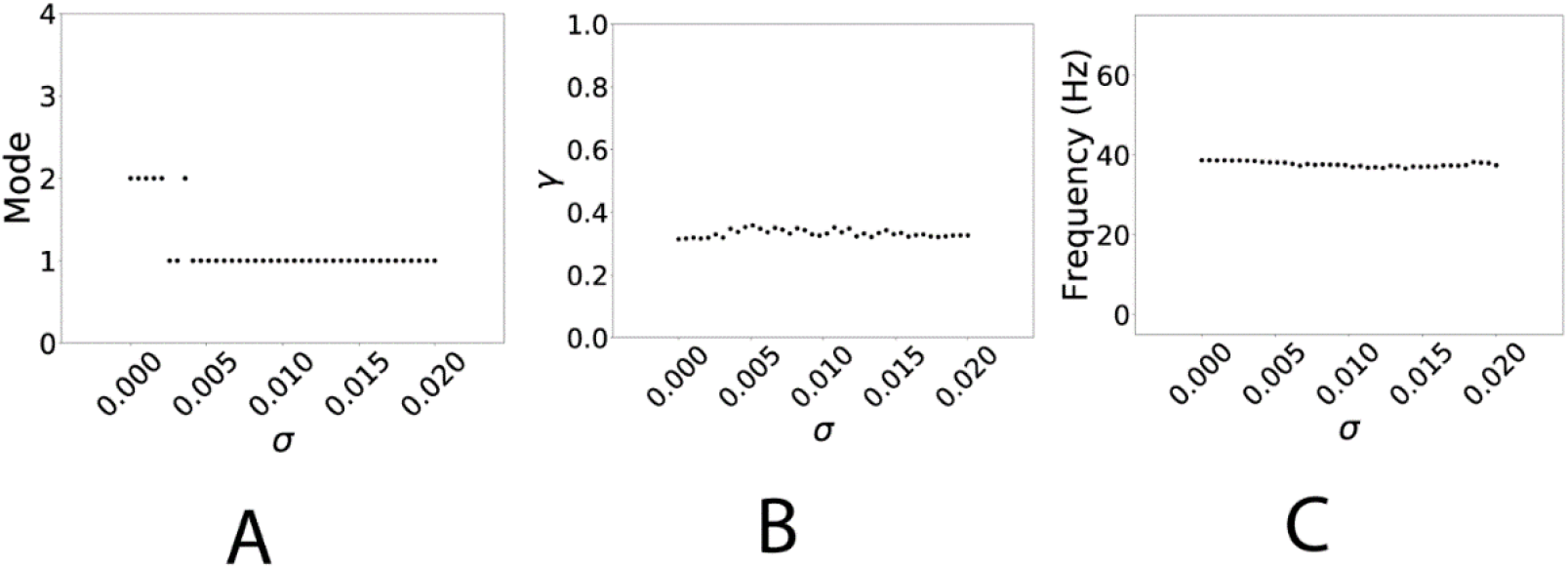
The effect of noise on synchrony properties of the network, which exhibits mode 2 desynchronization dynamics in the noiseless case (*ε*_1_ = 0.132). The strength of the noise *σ* is varied along the horizontal axes. A: mode of the system, B: average synchrony index γ, C: mean oscillation frequency (firing rate) of the system.

The effect of channel noise on the distribution of desynchronization durations for *ε*_1_ = 0.132 is shown in Figure 6. The strength of the noise increases from left to right, and the effect is to simultaneously broaden the distribution and shift the mode from 2 down to 1.

**Figure 6.**
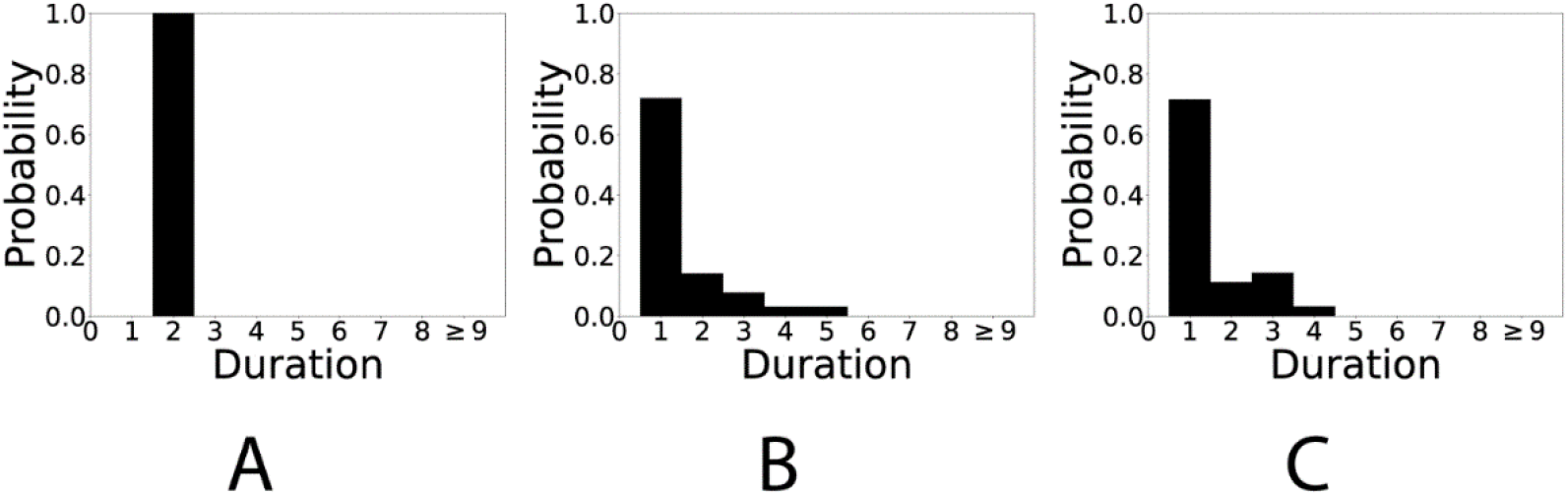
Distributions of desynchronization durations for different levels of noise in the network exhibiting mode 2 desynchronization dynamics in the noiseless case (*ε*_1_ = 0.132). Noise strength increases from left to right: A: *σ* = 0 (no noise), B: *σ* = 0.0097, C: *σ* = 0.02.

We note that the same trend was observed for systems with larger modes, i.e., a strong enough noise shifts the mode of a system down to one. For example, for *ε*_1_ = 0.184 the noiseless system is mode 4. However, if the noise is sufficiently large (yet small in a sense that it does not alter the shape of oscillations much and does not alter average synchrony), it will shift the system to mode 1 dynamics.

### 3.2. Variation of *β*

We again will first explore if the channel noise will preserve the prevalence of short desynchronizations or not under the variation of parameter *β*. The parameter *β* changes the widths of the voltage-dependent time-constant of activation *τ*(*v*) and the width of the steady-state activation function *w*_∞_(*v*) for potassium current. Large values of *β* correspond to the larger width of the sigmoidal function *w*_∞_(*v*) and to the larger width of the bell-shaped function *τ*(*v*). Both changes effectively delay the activation of the potassium current and are known to promote short desynchronization dynamics (Ahn and Rubchinsky, 2017). For *β* = 0.131, the noiseless system is mode 1. We see in Fig. 7A that the mode of the system is unchanged as noise is added and its strength is increased. The synchrony index and the frequency of the system are virtually unchanged from that of the noiseless system and are therefore not plotted.

**Figure 7.**
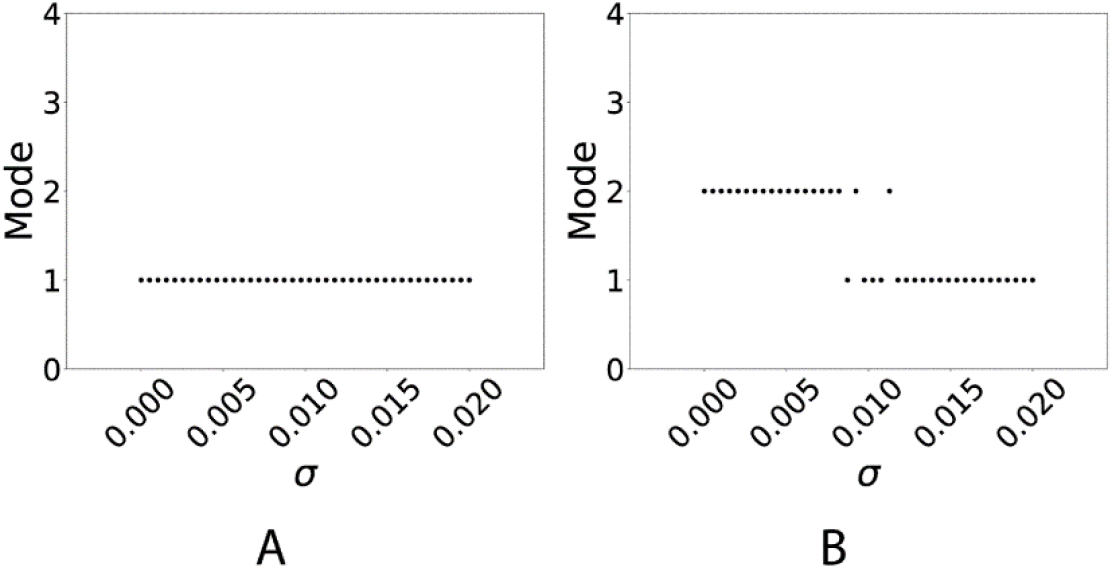
The effect of the noise strength *σ* on the mode of the desynchronization durations distribution for A: the network, which exhibits mode 1 desynchronization dynamics in the noiseless case (*β* = 0.131) and B: the network, which exhibits mode 2 desynchronization dynamics in the noiseless case (*β* = 0.080).

Now, let us look at a longer desynchronization case obtained by varying *β*. For *β* = 0.080, the noiseless system is mode 2. For small noise strengths, the system remains mode 2, but for larger values the system becomes mode 1 (Fig. 7B), although this transition is not fully monotonous with respect to the noise strength. Again, the shift in mode is independent of the average synchrony index or the mean firing frequency, which do not substantially change with noise (even though they are different from those in the network with *β* = 0.131).

### 3.3. Variation of *v*_*w*1_

The parameter *v*_*w*1_ affects a horizontal translation in *w*_∞_(*v*) and *τ*(*v*). In particular, increasing *v*_*w*_1 shifts both curves to the right, i.e., towards higher voltages. For the spike generation in the considered model neuron, it results in a faster potassium current activation. Smaller values of *v*_*w*1_ result in shorter desynchronization durations (Ahn and Rubchinsky, 2017). For *v*_*w*1_ = 0.096, the noiseless system is mode 1. All channel noise strengths considered preserve this mode 1 dynamics (Fig. 8A).

**Figure 8.**
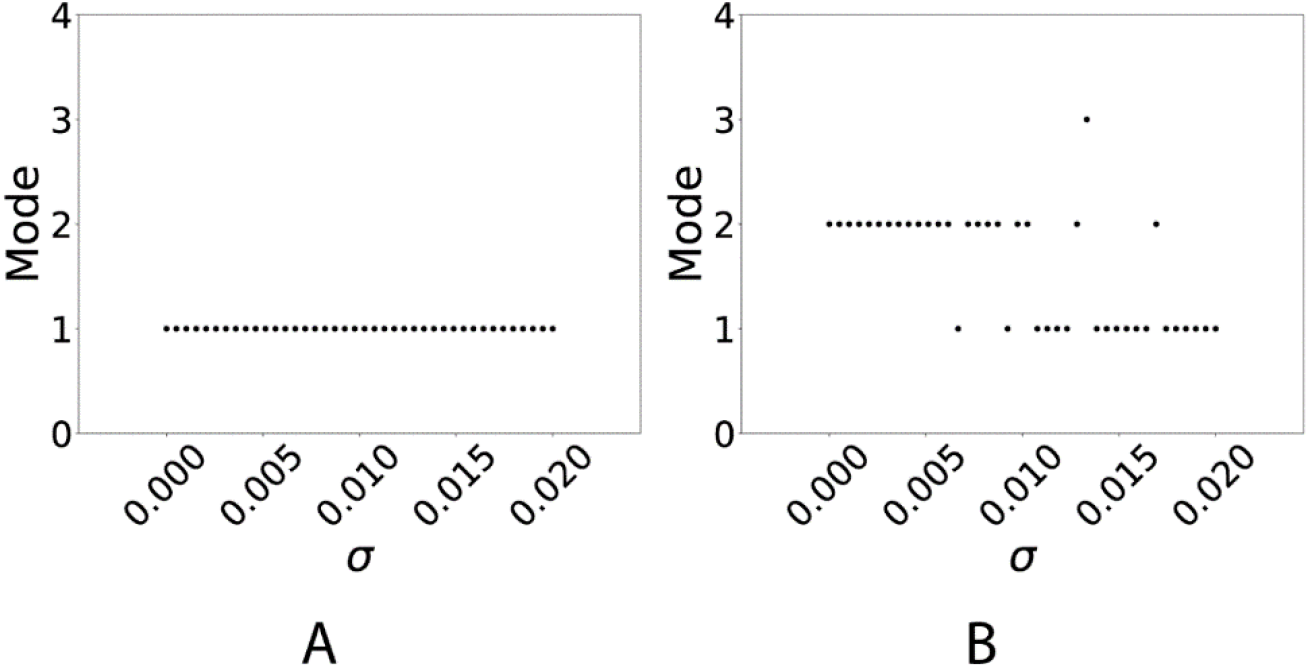
The effect of the noise strength *σ* on the mode of the desynchronization durations distribution for A: the network, which exhibits mode 1 desynchronization dynamics in the noiseless case (*v*_*w*1_ = 0.096) and B: the network, which exhibits mode 2 desynchronization dynamics in the noiseless case (*v*_*w*1_ = 0.169).

Now we set the parameter *v*_*w*1_ = 0.169 so as to place the noiseless network in mode 2 dynamics (longer desynchronizations). As we vary the noise magnitude from zero to larger values, the network exhibits a tendency for mode 1 (short desynchronizations) dynamics. This is not a monotonous transition, and with moderate noise the network may exhibit desynchronizations both shorter than in the noiseless case and longer than in the noiseless case. But a strong enough noise will shift the mode to one creating short desynchronization dynamics similar to experimentally observed ones (Fig. 8B). As earlier, in all the cases considered here, the frequency of oscillations is not altered, and the average synchrony strength only varies slightly (even though they are different in the network with different values of *v*_*w*1_ considered here).

### 3.4. Variation of *β_w_* and *β_τ_*

Variation of the previous parameters, i.e., *ε*, *β* and *v*_*w*1_, can affect the average synchronization strength and frequency of firing in addition to changing the durations of desynchronizations. For example, for *β* = 0.065 (and other parameter values as described above) the system exhibits a frequency of about 41 Hz (and desynchronization durations mode 1, 2 or 3 depending on the strength of the noise). While for *β* = 0.131 the frequency is about 14 Hz. Thus, when we are changing the values of parameters to explore the effect of noise on the temporal patterns of synchronization, we do not keep the frequency of oscillations and average synchrony strength fixed. To take care of this issue, i.e., to use noise to control the mode of a noiseless system while keeping both the average synchrony strength and firing frequency near constant, one can take the parameter *β* and separate it into two independent parameters, *β*_*τ*_ and *β*_*w*_, for *τ*(*v*) and *w*_∞_(*v*) respectively. The result is that the mode of the system is essentially independent of the synchrony strength and frequency as these two parameters are varied simultaneously (Ahn and Rubchinsky, 2017). A smaller *β*_*w*_ and larger *β*_*τ*_ result in shorter desynchronization durations even though frequency and average synchrony stay basically the same.

For *β*_*w*_ = 0.098, *β*_*τ*_ = 0.079 the noiseless system is mode 1. As illustrated in Fig. 9A1, the mode remains at one even with the introduction of noise, regardless of the noise strength.

**Figure 9.**
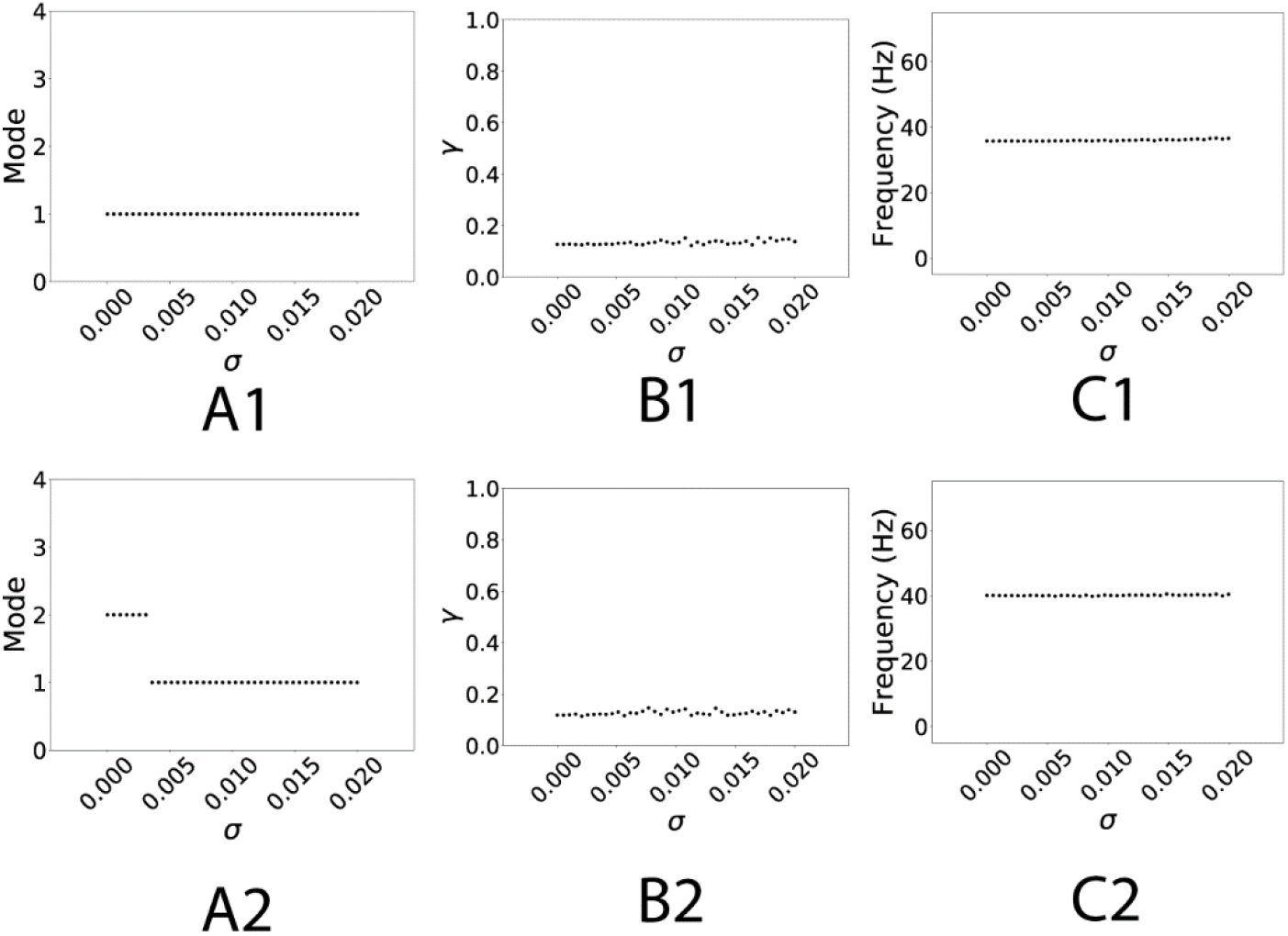
The effect of the noise on the mode of the desynchronization durations distribution in the networks, which exhibits mode 1 desynchronization dynamics (row 1, *β*_*w*_ = 0.098, *β*_*τ*_ = 0.079) or mode 2 desynchronization dynamics (row 2, *β*_*w*_ = 0.120, *β*_*τ*_ = 0.068) in the noiseless case, but otherwise have similar average synchronization strength and firing rate. The strength of the noise *σ* is varied along the horizontal axes. A: mode of the distribution of desynchronization durations, B: average synchrony index γ, C: mean oscillations frequency (firing rate) of the system.

For *β*_*w*_ = 0.120, *β*_*τ*_ = 0.068 the noiseless system is mode 2, moreover, the average frequency of oscillations and the average synchronization strength stay the same as in the mode 1 case above. Introduction of noise of sufficiently large strength again leads to shortening of desynchronizations and mode 1 dynamics, see Fig. 9A2. Not only do the frequency and average synchrony strength not change with respect to noise (Fig. 9B2 and 9C2), but their values are not substantially different from those in Fig. 9B1 and 9C1.

### 3.5. Additive Noise

We also consider the action of an additive noise (current noise) on the temporal patterning of the synchronized dynamics. Thus, we consider here the action of additive (current, activity-independent) noise as opposed to the multiplicative (channel, activity-dependent) noise. We use the same framework, looking at different variations of parameters and resulting modes of desynchronization duration distributions as described in the previous sections. Since the effects of parameter selection has been already described above, we will just briefly present the results here.

In general, the effect of an additive noise on a system is very similar to the effect of the conductance noise, i.e., the mode of a system is switched down to one at sufficiently strong noise strengths. Thus, the forthcoming Fig. 10 is similar to figures in 3.1-3.4 above. The changes of average synchrony and firing frequency are not plotted, because they do not experience any substantial variation for the considered range of noise strength. For variation of *ε* results refer to the first row in Fig. 10, for variation of *β* results refer to the second row in Fig. 13, for variation of *v*_*w*_1 results refer to third row in Fig. 10, and for variation of *β*_*w*_, *β*_*τ*_ results refer to the last row in Fig. 10.

**Figure 10.**
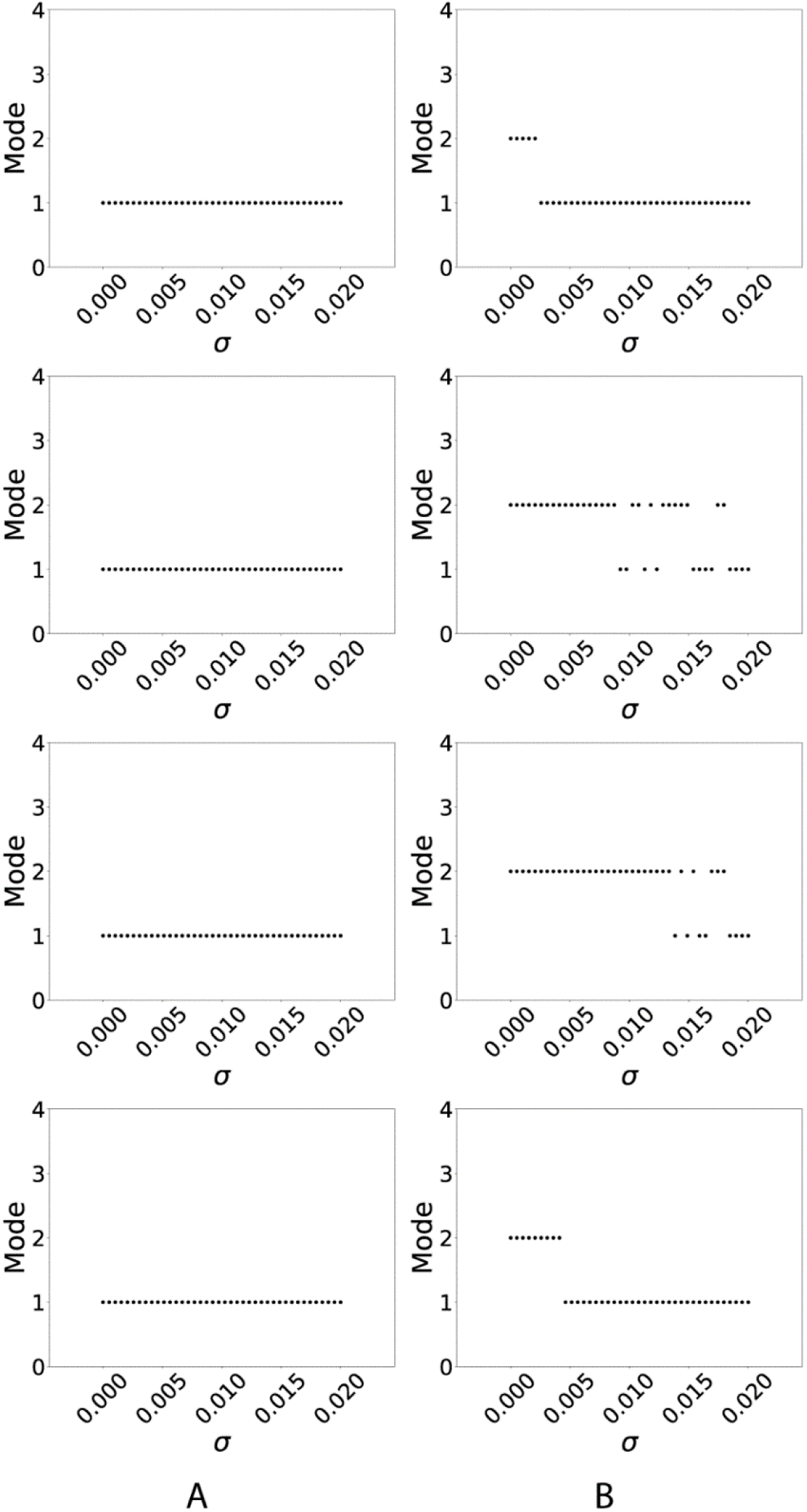
The effect of additive noise on mode of the desynchronization durations distribution. The strength of the noise is varied along the horizontal axes. A: the network, which exhibits mode 1 desynchronization dynamics in the noiseless case, B: the network, which exhibits mode 2 desynchronization dynamics in the noiseless case. The four rows present the networks with parameters considered in the sections 3.1-3.4, except that the noise is additive here.

## 4. DISCUSSION

Our study considered intermittent synchronization of oscillations in a minimal network of two synaptically coupled neurons. In general, as the coupling strength between oscillators increases, the synchronization strength will increase from a low to high level (Pikovsky et al., 2001). Moderate values of coupling naturally give rise to intermittent synchrony, where episodes of strong synchrony are interspersed with episodes of desynchronized dynamics. The same level of synchrony may be reached with a few long desynchronizations, many short desynchronizations, and many possibilities in between. Here we considered how noise will affect the temporal patterning of moderately synchronous dynamics.

We showed that weak noise may alter the temporal patterning of synchrony and starting from certain magnitude it leads to the synchronized activity punctuated by very short desynchronizations. That is, with noise of sufficient magnitude, the distribution of desynchronization durations has a mode of one. This is the case for both channel (or conductance) noise and current noise (i.e., for both multiplicative and additive noise). Interestingly, this reorganization of the temporal patterning of synchrony is achieved with relatively weak noise, so that the average synchronization strength is not substantially changed.

The observed phenomena are interesting to consider in the context of the experimental results on the temporal patterning of neural synchrony in the brain. Multiple experiments show that moderately synchronized brain activity at rest has a very specific temporal pattern: most of the desynchronizations are very short, lasting for just one cycle of oscillation (Park et al., 2010; Ahn and Rubchinsky, 2013; Ahn et al., 2014; Malaia et al., 2020; see also Introduction). This study suggests that noise may be one of the factors promoting this regime of desynchronization dynamics. Naturally, real neurons are noisy, and there may be many sources of this noise (in particular, depending on the scale of the system considered). Multiplicative noise considered here is a good way to describe inherent stochasticity of ion channels in neural membranes (Goldwyn and Shea-Brown, 2011). Additive noise may represent stochastic inputs to neurons. Both are naturally occurring in the brain and the effect of either on the temporal patterning of synchrony is robust: it promotes short desynchronizations as in the experiments.

Other potential mechanisms of experimentally relevant short desynchronizations dynamics have been considered and include the specific kinetics of the ionic channels (Ahn and Rubchinsky, 2017) and spike-timing dependent plasticity (Zirkle and Rubchinsky, 2020). The stochastic mechanism of the short desynchronization dynamics considered here is not necessarily mutually exclusive with those mechanisms, rather all of them may potentially act in a cooperative manner.

It is important to note the limitations of the study. The network studied here is a minimal network. Real neural networks of the brain have, of course, a much more complicated organization. The same consideration applies to the relatively simplistic model neurons used here. However, the robustness of the effect of noise even in this simple system may indicate that it is likely to persist on a larger scale. Also, this study does not provide an exhaustive quantitative description of the noise action. Rather it illustrates what the noise is capable of in terms of synchronization dynamics changes.

Finally, we would like to note that noise may exert different positive effects in the brain, such as increased reliability, sensitivity, and regularity (e.g., Ermentrout et al., 2008; Faisal et al., 2008). For example, a well-known effect of stochastic resonance (response to a weak signal improved by noise) has been described in neuronal networks and the whole brain experimentally (e.g., Gluckman et al. 1996; Ward et al., 2010). It also has long been discussed that noise (and, broadly speaking, various irregularities) may benefit neural systems because irregularities help networks to exhibit a wide repertoire of different dynamics (e.g., Rabinovich and Abarbanel, 1998; Ghosh et al., 2008; Garrett et al., 2011; McDonnell and Ward, 2011; Yarom and Hounsgaard, 2011). Perhaps somewhat similarly, noise effects the temporal variability of neural synchrony in such a way as to create a system with very dynamic behavior in the form of short desynchronizations dynamic. The latter may (as was argued in Ahn and Rubchinsky, 2013, 2017) lead to a network which can quickly, and efficiently, form and break-up transient neural assemblies to perform associated neural functions.

## Acknowledgements

This work was supported by NSF DMS 1813819.

